# Cryo-electron tomography reveals that dynactin recruits a team of dyneins for processive motility

**DOI:** 10.1101/182576

**Authors:** Danielle A. Grotjahn, Saikat Chowdhury, Yiru Xu, Richard J. McKenney, Trina A. Schroer, Gabriel C. Lander

## Abstract

A key player in the intracellular trafficking network is cytoplasmic dynein, a protein complex that transports molecular cargo along microtubule tracks. It has been shown that vertebrate dynein’s movement becomes strikingly enhanced upon interacting with a cofactor named dynactin and one of several cargo-adapters, such as BicaudalD2. However, the mechanisms responsible for this increase in transport efficiency are not well understood, largely due to a lack of structural information. We used cryo-electron tomography to visualize the first 3-dimensional structure of the intact dynein-dynactin complex bound to microtubules. Our structure reveals that the dynactin-cargo-adapter complex recruits and binds to two dimeric cytoplasmic dyneins. Interestingly, the dynein motor organization closely resembles that of axonemal dynein, suggesting that cytoplasmic dynein and axonemal dyneins may utilize similar mechanisms to coordinate multiple motors. We propose that grouping dyneins onto a single dynactin scaffold promotes collective force production as well as unidirectional processive motility. These findings provide a structural platform that facilitates a deeper biochemical and biophysical understanding of dynein regulation and cellular transport.

## Introduction

Precise spatial and temporal delivery of components to specific locations within a cell requires tightly regulated trafficking across a vast microtubule (MT) network (Welte 2004). A key player in intracellular trafficking is cytoplasmic dynein-1 (hereafter dynein), which transports molecular cargo towards MT minus ends. Dynein functions as a multi-subunit complex of dimerized “heavy chains” (DHCs), containing a carboxy-(C)-terminal “motor” domain and an amino-(N)-terminal “tail” region that contains a dimerization domain and attachment sites for several non-catalytic subunits. The dynein motor is distinct from other cytoskeletal motors, composed of an AAA+ ATPase ring interrupted by a coiled-coil stalk with a globular microtubule-binding domain (MTBD) (Schmidt and Carter 2016, Zhang 2017). Notably, purified vertebrate dynein exhibits limited, diffusive movement on MTs. Long-range, minus end-directed movement requires the association of dynactin, a megadalton-sized multi-subunit cofactor, as well as one of various cargo adaptors, such as the N-terminal fragment of BicaudalD2 (BICD2N) (McKenney 2014, Schlager 2014). Mutations that disrupt these dynein-cofactor interactions are associated with a variety of neurological pathologies (Hoang 2017). Although the manner by which BICD2N structurally mediates interactions between the dynein tail and dynactin has been elucidated by cryo-EM (Urnavicius 2015), a fundamental question remains: How do interactions with the dynein tail confer unidirectional processivity on the dynein motor domains?

### Structure determination of microtubule-bound dynein-dynactin-BICD2 complex

To understand how dynein is harnessed to yield processive movement, we isolated mouse dynein-dynactin-BICD2N (DDB) complexes bound to microtubules following methods previously described (Chowdhury 2015), and used cryo-ET and subtomogram averaging to determine the 3D structure (Figure 1, Supplemental Figure 1, Supplemental Movie 1). To facilitate the 3D reconstruction of this inherently heterogeneous complex, we incorporated an assisted alignment procedure into the RELION subtomogram averaging workflow(Bharat and Scheres 2016), followed by focused refinement of individual components (dynein tails-dynactin-BICD2N (TDB), and each pair of dynein motors) (Supplemental Figures 2 and 3, see Methods). The resulting structures were merged in UCSF Chimera (Goddard 2007) to obtain the final reconstruction of the intact DDB-MT complex (Figure 1 and Supplemental Figure 4, see Methods).

**Figure 1.**
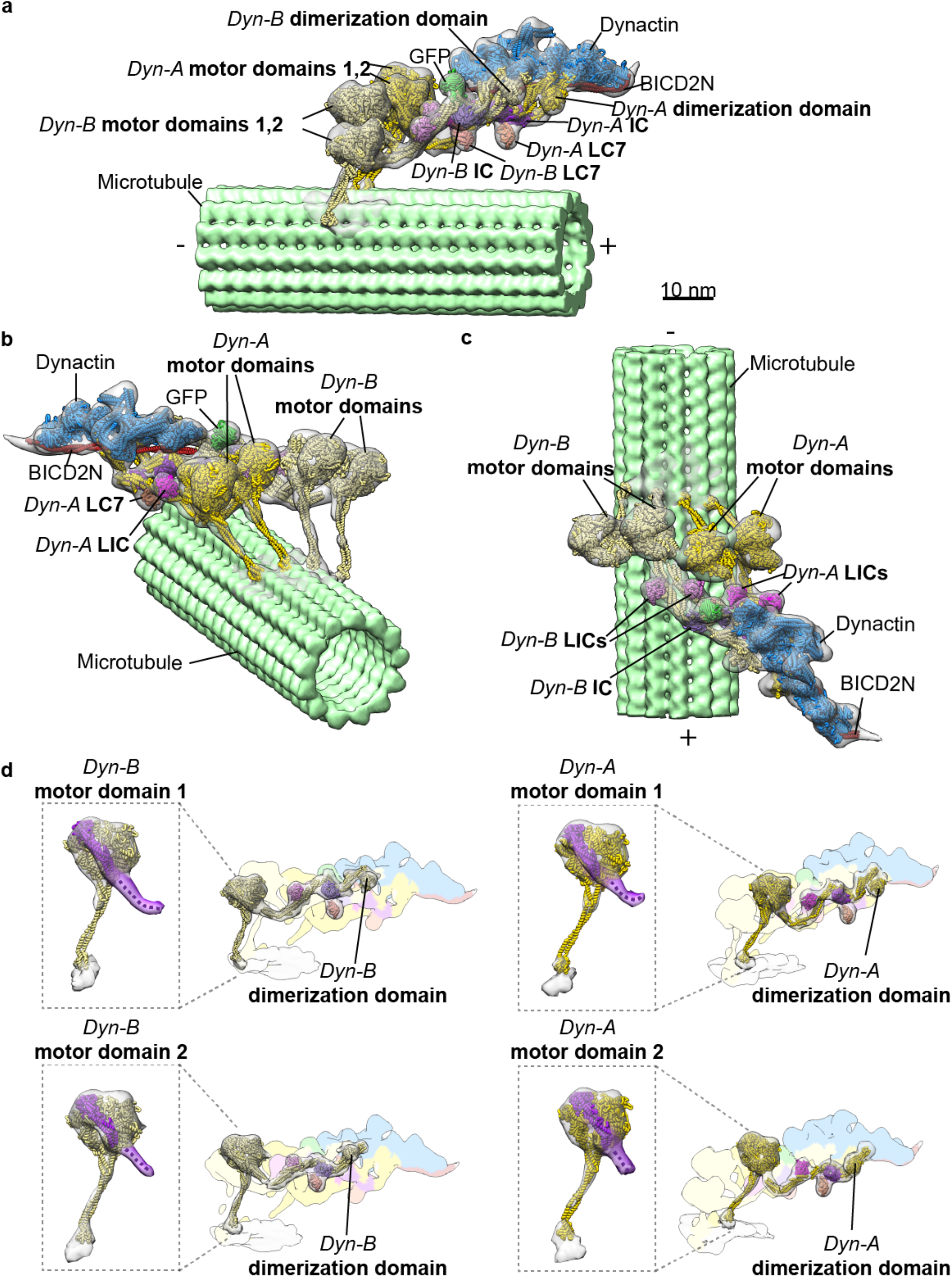
3D organization of the microtubule-bound dynein-dynactin-BICD2N complex. **a-c**, Three views of the subtomogram average (gray transparent density) of the MT-DDB complex are shown, with fitted atomic models of dynein dimer-1 (Dyn-A; yellow), dynein dimer-2 (Dyn-B; light yellow), dynactin (blue), BICD2N (red), associated chains (purple, salmon, magenta), and the BICD2N GFP tag (green), and a microtubule model (light green) PDBs used in fitting listed in Table 1. **d**, Cryo-EM density of each dynein motor domain (boxed region) shows the linker arm (purple) in the post-powerstroke conformation, consistent with AMPPNP binding. Cryo-EM density for each dynein HC and associated subunits with docked models, with the remainder of the cryo-EM density colored according to component composition (all coloring as in a-c).

### BICD2N mediates the association of two dynein dimers with a single dynactin

The overall organization of the DDB-MT resembles previous structures (Chowdhury 2015, Urnavicius 2015), but a striking new feature emerged: the presence of two complete dimeric dynein densities bound to dynactin (Figure 1). The details of the reconstruction were sufficient to visualize the entirety of the four DHCs from the dynactin-bound N-terminus to the C-terminal motor domains, and to confirm the post-power-stroke conformation of the motor linker domain (Schmidt 2012, Bhabha 2014), which is consistent with the presence of AMPPNP during the isolation procedure (Figure 1d). The four motor domains are positioned in a row, ~17nm from the MT surface, with weak density attributable to the stalk contacting the corresponding MT. Additionally, the structure displays densities for several other dynein subunits, including the light intermediate chain (LIC), light chain 7 (LC7), and intermediate chain (IC) (Figure 1), in positions that are consistent with previous studies (Chowdhury 2015, Urnavicius 2015, Zhang 2017). By fitting available atomic models into the EM density, we generated a pseudo-atomic model of the complete DDB-MT complex, including two dynein dimers (Dyn-A and Dyn-B) with associated dynein subunits, one dynactin-BICD2N complex, as well as the GFP-tag at the N-terminus of BICD2N.

Manual inspection of the raw extracted subtomograms revealed that over 97% of the dynactin densities were associated with four dynein motor domains (Figure 2a, Supplemental Figure 1b). Importantly, a focused 3D classification on the region surrounding the dynein motor domains did not yield any well-resolved 3D classes containing a single dynein dimer (Supplemental Figure 5b), reinforcing our conclusion. Furthermore, comparison of our reconstruction with previously determined 2D averages of negatively stained DDB-MT complexes (Chowdhury 2015) revealed highly correlated structural features (Supplemental Figure 5c), suggesting that two dynein dimers were associated with a single dynactin in our earlier 2D averages of DDB-MT complex (Chowdhury 2015), but the limitations of 2D negative stain precluded resolution of multiple dynein motors in that study. Overall, these results strongly support a model in which BICD2N can facilitate binding of two dynein dimers to a single dynactin complex.

**Figure 2.**
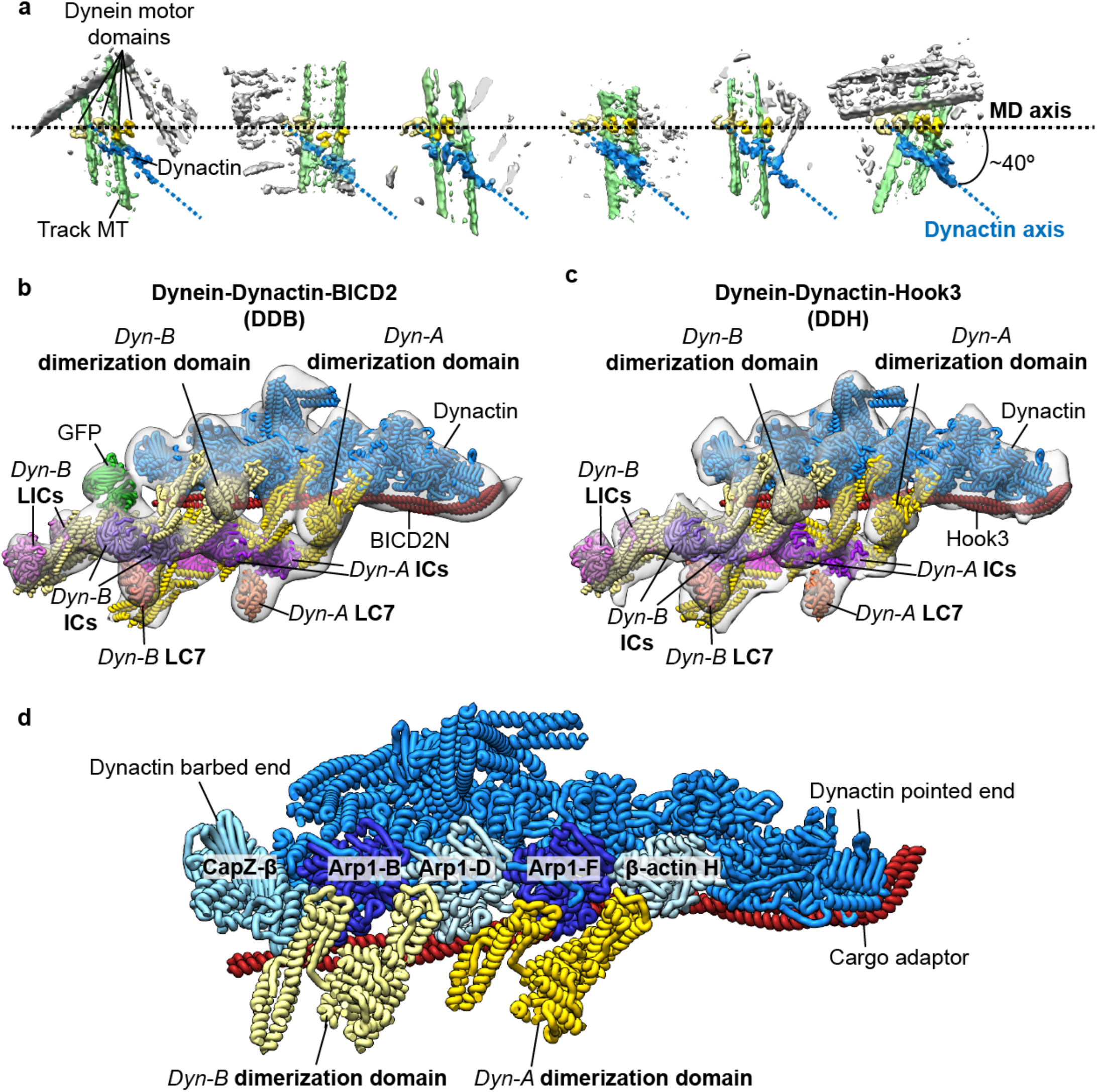
Association of two dyneins with dynactin in the presence of cargo adaptor proteins. **a**. Raw subtomograms show that dynein dimers (motor domains (MDs) colored in two shades of yellow) associate with a single dynactin (blue) in Dyn-adaptor-MT complexes. The MDs are arranged horizontally (axis represented by black dotted line) showing that the dynactin is oriented at a ~40° relative to the MD to the axis. The DDB-associated MT is colored green, non-associated MTs are colored gray. **b-c**. Subtomogram averages (gray transparent density) of the dynactin-dynein tail-cargo adaptor portion of the DDH-MT (a) and DDB-MT (b) complexes with docked atomic models of dynein tails (colored as in Figure 1). Both complexes present a similar overall architecture with two dimeric dyneins bound to a single dynactin. **d**. A pseudo-atomic model of the dynactin-dynein tail-cargo adaptor complex shows interactions between two dimeric dynein tails and the dynactin filament. The tail of Dyn-A binds to dynactin across Arp1-F subunit with one heavy chain binding at the interface between ß-actin H and Arp1-F, and the other chain binds at the interface between Arp1-F and Arp1-D. The tail of Dyn-B binds across Arp1-B subunit of dynactin with one heavy chain binding at the interface between Arp1-B and D subunits and the other between Arp1-B and CapZ-β.

The observation that the dynactin-BICD2N assembly binds to two dynein dimers in the presence of MTs is unexpected because prior motility assays and structural studies concluded that only one dynein dimer was present in the dynactin-BICD2N complex (McKenney 2014, Schlager 2014, Urnavicius 2015, Zhang 2017) (Figure 1, Supplemental Figure 5a,c). However, prior motility assays reported that a subset of DDB complexes exhibited extreme run lengths (>50 μm) (McKenney 2014), which might be attributed to DDB complexes containing two dynein dimers. Recent single molecule experiments show that DDB complex velocities on MTs distribute into two populations, with one exhibiting twice the velocity of the other (Gutierrez 2017). Additionally, recent structural studies have shown that dynactin-BICD2N is capable of binding two dimeric dyneins (A. P. Carter, personal communication). It is possible that inclusion of AMPPNP in the lysate, which immobilizes DDB complexes on MTs for structural analyses, induces a dynein conformation that favors attachment of two dynein dimers to dynactin-BICD2. Together, our data suggest that regulatory mechanisms exist that influence the DDB’s dynein:dynactin stoichiometry.

### Hook3 also recruits two dynein dimers to dynactin

To assess whether the recruitment of two dynein dimers is unique to the BICD2N scaffold, we isolated dynein-dynactin complexes bound to MTs in the presence of another cargo adaptor, an N-terminal fragment of Hook3, which was also shown to endow dynein-dynactin with processive motility (McKenney 2014, Olenick 2016, Schroeder and Vale 2016). Strikingly, the subtomogram average of the resulting dynein-dynactin-Hook3 (DDH) complex again revealed two tail domains interacting with dynactin and EM density attributable to two sets of dynein’s accessory subunits (LC, IC, LIC) (Figure 2b, Supplemental Figure 6). The fact that the structures of the DDH and DDB are largely indistinguishable (Figure 2b, c, Supplemental Figure 6) suggests that recruitment of two dynein molecules to the dynactin-cargo adaptor complex is a widely conserved mechanism for inducing processive motility.

### Dynein tails bind to adjacent clefts on the Arp1 filament of dynactin

Our 3D reconstruction illustrates how one dynactin-adaptor complex can accommodate two dynein dimers. The previously determined TDB structure showed the dynein tail to be bound to two clefts along dynactin’s Arp filament: one between Arp1-D and F, and the other between Arp1-F and β-actin H (Urnavicius 2015). We observe identical interactions here (Fig 2b, c). The second dynein tail binds the Arp1 filament in a highly similar fashion, interacting with two adjacent clefts near the barbed-end of dynactin, one between the Arp1-D and B and the other between Arp1-B and the CapZ-dimer (Fig 2 b, c, d). The fact that neither our study, nor previous studies, observe complexes in which dynein straddles the clefts in the center of the Arp filament (i.e., on either side of Arp1-D) suggests that the dynactin-cargo adaptor interface has evolved to maximize dynein occupancy on dynactin.

### Motor domains are positioned for processive motility

In contrast to previous structural studies of isolated DDB complexes (Chowdhury 2015, Urnavicius 2015), our structure reveals the spatial organization of the dynein motor domains (MDs) relative to the dynactin complex. The four MDs are equidistantly spaced ~12 nm apart, with all four MTBDs projecting towards the MT minus end (Figure 1 b, c, Supplemental Figure 5d). Interestingly, there is some variability in the transverse angle at which the MD pairs attach to the MT axis, which limits our ability to resolve individual tubulin dimers in the MT lattice (Figure 2a). Regardless, the spacing between the MTBDs is consistent with the MT helical protofilament spacing, suggesting that the four MDs associate with four distinct but adjacent MT protofilaments (Supplemental Figure 5d). Notably, interactions of the dynein tail with dynactin’s helical Arp filament yield a conspicuous “skewed” organization in which dynactin is oriented approximately 40º relative to the linear array of dynein motors (Figure 2a).

To confirm that the dynein MD configuration on MTs is promoted by the dynactin-adaptor complex, we used cryo-ET to visualize dynein dimers bound to MTs in the absence of dyanctin and adaptors. Manual inspection of 229 sub-volumes showed that isolated dynein dimers bind the MT surface individually, with their motor domains at a range of distances from one another (Supplemental Figure 7), hindering our ability to generate a 3D average of these complexes. Despite this complication, our results suggest that in the absence of cofactors, individual cytoplasmic dynein complexes bind individually to the MT, with the two MDs positioned at variable distances from one another. Thus, not only does the dynactin-cargo adaptor complex recruit multiple dyneins, it positions their MDs in an array highly compatible with unidirectional processive movement. This is consistent with prior work showing that association of a single dynein with dynactin results in a dramatic reorganization of dynein from an auto-inhibited conformation to one that is capable of productive minus-end movement (Zhang 2017).

### Structural and mechanistic parallels between cytoplasmic and axonemal dynein

In addition to positioning the dynein MDs for processive motility, dynactin can also serve as a scaffold for collective force production. Vertebrate dynein motors have been shown to work collectively to generate forces that far exceed those produced by an individual dynein motor (Rai 2013, Rai 2016), and this multi-motor coordination may be required to carry out high-load transport processes, such as nuclear positioning, mitotic spindle rotation, and organelle trafficking. A well-characterized example of teamwork among dynein motors can be found in cilliary and flagellar axonemes, where axonemal dyneins are known to work in huge ensembles to accomplish large-scale, synchronized, cilliary and flagellar motility (Wemmer and Marshall 2004).

We wondered if the dynein configuration observed in our structures showed any similarities to axonemal dynein. In addition to working collectively in teams to accomplish large-scale, synchronized, cilliary and flagellar motility (Wemmer and Marshall 2004), axonemal dyneins contain a C-terminal motor domain that is similar to cytoplasmic dynein, even though they have evolved a distinct N-terminal tail to accommodate its cellular function (Ishikawa 2012). Intriguingly, the organization of the dynein motors in the DDB-MT structure is strikingly similar to that of sea urchin sperm flagella outer dynein arms in the post-powerstroke state (Figure 3a, Supplemental Figure 8) (Lin 2014). In both structures, the dynein tails are associated with an elongated, filamentous structure – a microtubule doublet in the case of axonemal dynein, and dynactin’s actin-like filament in the case of cytoplasmic dynein (Figure 3b). This leads to the intriguing hypothesis that cytoplasmic and axonemal dyneins utilize a similar mechanism for coordinating the activity of multiple dynein motors, in which parallel arrangement of the motor domains relative to the MT allows the conformational change associated with ATP hydrolysis to propel the MTBD more effectively toward the MT minus end (Figure 3c). By contrast, ATP hydrolysis in the absence of the massive dynactin scaffold might result in non-directed motion of the stalk or unproductive motion of the dynein tail, limiting the propensity for minus end movement (Figure 3c). Another non-mutually exclusive possibility is that the second motor might increase the duty ratio of the entire complex by providing an additional attachment to the MT lattice and reducing the probability of complex dissociation from the MT during movement. Such an effect could enhance complex processivity, as has been observed in assays that multimerize motors on an artificial scaffolding, such as a bead or DNA chassis (Derr 2012, Xu 2012, Torisawa 2014).

**Figure 3.**
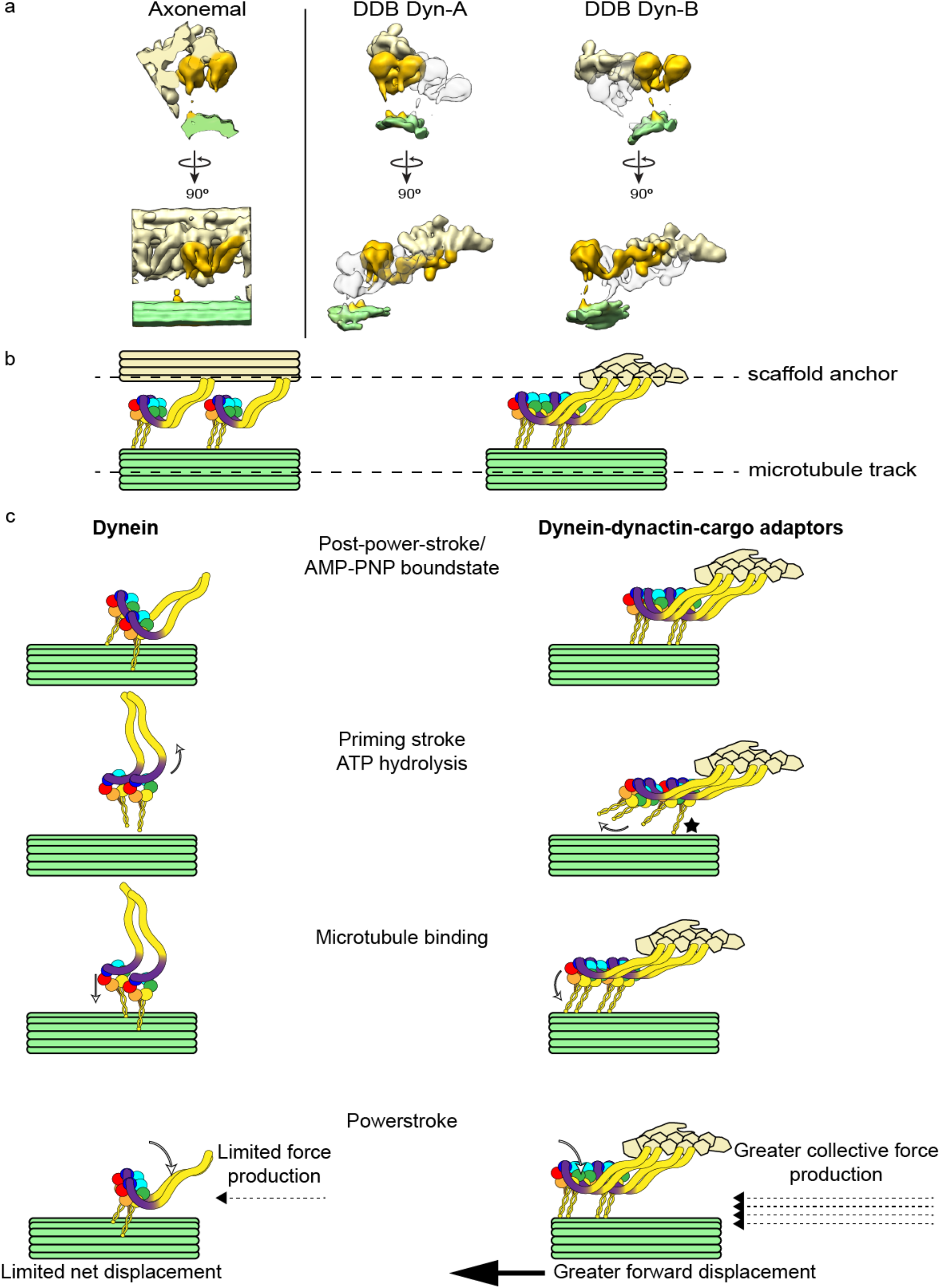
Organizational and mechanistic commonalities between axonemal dynein and cytoplasmic dynein, suggesting a model for processivity. **a**. Orthogonal views of the axonemal dynein subtomogram average (EMD-5757 (Lin 2014)) are shown in the left panel. Axonemal dynein (golden) associates with a MT doublet scaffold (light yellow) through its tail and another MT doublet (green) through the MT binding stalk of the motor. The right side panel shows the organization of cytoplasmic dyneins in dynein-dynactin-cargo adaptor-MT complexes. Each of the two dimeric dyneins (Dyn-A and Dyn-B) are highlighted in gold and associate with the dynactin scaffold (light yellow) via the tails and to MT surface (light green) through the MT binding stalk of the motors. **b**. Similarities between the overall organization of multiple axonemal dyneins in axoneme (left) and two cytoplasmic dyneins in Dyn-cargo adaptor-MT complexes (right) are shown using diagrammatic representations. Each AAA+ domain with the dynein motor domain is colored uniquely, with linker arm colored purple. In both systems, multiple dyneins are associated with a filamentous scaffold (a MT doublet or dynactin) via N-terminal tail interactions. The dynein motors associate with MT tracks through the binding stalk. In this way, both axonemal and cytoplasmic dyneins integrate into scaffolds to work in teams. c. In Dyn-adaptor-MT complexes (right column), by anchoring the dynein tail to a scaffolding structure, ATP hydrolysis results in a rotation that propels the MTBD towards the MT minus end, positioning the MT binding stalk further along the MT upon its return to a high affinity state. The dynein powerstroke thus results in productive translocation of the scaffold and attached cargo toward the MT minus end. Star denotes MT-interacting stalk that prevents dissociation of the complex from the MT. In contrast, in absence of a scaffold such as dynactin (left column), there is an increased likelihood that ATP hydrolysis results in unproductive motion of the dynein tail, limiting the propensity for minus end movement. Additionally, association of multiple dyneins with a dynactin scaffold results a reduced probability of complex dissociation during processive movement, and in larger collective force production, similar to axonemal dyneins.

The collective force generation previously shown for cytoplasmic dynein (Rai 2013) may also involve an element of synchronization, as observed in axonemal dynein (Lin 2014). It has been proposed that axonemal dynein motors synchronize as coupled oscillators by virtue of their connection to a shared scaffold (Wemmer and Marshall 2004). Our reconstructions show that dynactin is a scaffold for cytoplasmic dynein that may promote similar synchronization among bound dynein motors. Motor synchronization may also involve physical contacts between adjacent motor domains to promote teamwork and collective force generation. Interestingly, we observe density between MDs extending from AAA3 domain to the linker domain of the neighboring MD in the DDB reconstruction (Supplemental Figure 9). It is possible that this density corresponds to an inter-MD interaction involving the DHC C-terminal extension (CT-cap), which has been shown to play a role in regulating dynein processivity in Dictyostelium (Numata 2011) and vertebrates (Nicholas 2015). It is also possible that this bridging density corresponds to an auxiliary regulator factor, such as Lissencephaly-1, which binds directly to the dynein MD (Toropova 2014, DeSantis 2017) and has been shown to increase velocity of vertebrate DDB complexes (Baumbach 2017, Gutierrez 2017). Higher resolution studies will be required to confirm these hypotheses.

In conclusion, we used cryo-ET and subtomogram averaging to reveal the 3-dimensional structure of intact dynein-dynactin-cargo adapter complexes bound to MTs, and discovered that multiple distinct cargo adapters are able to mediate the association of two dynein dimers with a single dynactin. This novel configuration imposes spatial and conformational constraints on both dynein dimers, positioning the four dynein MDs in close proximity to one another and oriented towards the MT minus end. Grouping multiple dyneins onto a single dynactin scaffold has the potential to promote collective force production, increased processivity, and favor unidirectional movement, suggesting mechanistic parallels to axonemal dynein. These findings provide a platform that integrates decades of biochemical and biophysical studies on the unusual behavior of this large, highly conserved, minus end-directed motor protein, while posing further interesting questions regarding the underlying mechanisms of dynein-mediated intracellular transport.

## Acknowledgements

We thank Jean-Christophe Ducom at The Scripps Research Institute High Performance Computing for computational support, and Bill Anderson at The Scripps Research Institute electron microscopy facility for microscope support. We also thank Elliot Mattson for input on the manuscript. D.G. is supported by a National Sciences Foundation predoctoral fellowship. G.C.L. is supported as a Searle Scholar, a Pew Scholar, and by the National Institutes of Health (NIH) DP2EB020402. R.M is supported by the National Institutes of Health (NIH) R00 grant R00NS089428. T.S. is supported by the Johns Hopkins Krieger School of Arts and Sciences. Computational analyses of EM data were performed using shared instrumentation funded by NIH S10OD021634 to G.C.L.

## Methods

### Purification of MT-bound complexes

The cargo adaptor proteins GFP-BICD2N (a.a. 25-400) and SNAPf-Hook3 (a.a. 1-552) were recombinantly expressed and purified as described previously (McKenney 2014, Chowdhury 2015). MT-bound DDB complexes were prepared from mouse brain tissue as described previously (Chowdhury 2015). Isolation of MT-bound DDH complexes was performed using the same MT-DDB protocol, with a minor modification to incorporate aspects of a protocol established by (Amos 1989) to enrich DDH complex on MTs. We initially removed bulk tubulin from the lysate by adding 6 μM Taxol and 0.2 mM GTP, performing one round of MT polymerization, and then pelleting and discarding the polymerized MTs and MAPs by centrifugation. In order to prevent endogenous dynein from associating with the MTs prior to pelleting, 0.5mM Mg^2+^-ATP was added to the lysate. This resulted in lysate having a higher dynein-to-tubulin ratio. The remaining tubulin in lysate was then polymerized by adding 10 μM Taxol and 1 mM GTP, and 4mM Mg^2+^ - AMPPNP and 500nM of Hook3 was added to promote engagement of the DDH complexes to the MTs.

MT-engaged dynein was prepared from mouse brain using similar procedures as described for MT-DDH complex, but to prevent the association of endogenous dynactin with dynein, the lysate was not supplemented with recombinant cargo adaptor proteins. The protocol for this work was approved by the TSRI IACUC office under protocol 14-0013.

### Grid preparation for cryo-EM analysis

All samples were prepared for cryo imaging in a similar manner. The complex-bound MT pellets were diluted 20 fold with PMEE buffer supplemented with 1mM GTP, 4mM Mg^2+^ - AMPPNP, and 20μM Taxol at room temperature. 5nm colloidal gold (Ted Pella) were pretreated with BSA to prevent aggregation as described previously (Iancu 2006). Immediately before freezing, samples were diluted 60 to 120-fold and mixed with the pre-treated colloidal gold (optimal dilution for each sample was determined by screening the cryo-EM grids at a range of concentrations). 4 μl aliquots of sample were applied to freshly plasma-cleaned (75% argon / 25% oxygen mixture) UltrAuFoil grids (Quantifoil) containing holes 1.2 μm in diameter spaced 1.3 μm apart. Plunge freezing was performed using a custom-built manual plunging device. The grid was manually blotted from the side opposite to which the sample was applied with a Whatman 1 filter paper for 5-7 s to remove excess sample. After blotting, the grid with remaining sample was immediately vitrified by plunge freezing into a liquid-ethane slurry. The entire procedure was carried out in a cold room maintained at 4°C and >90% relative humidity.

### Cryo-ET data acquisition

Tilt series for DDB-MT and DDH-MT samples were collected using a Thermo Fisher Titan Krios electron microscope operating at 300 keV and equipped with a Gatan K2 Summit direct electron detector. Data acquisition was performed using the UCSF tomography package (Suloway 2009) implemented within the Leginon automated data acquisition software (Suloway 2005). Tilt series were acquired using a sequential tilting scheme, starting at 0° and increasing to +59º at 1° increments, then returning to 0º and increasing to -59º at 1° increments. Each tilt series was collected with a nominal defocus value that was randomly set to between 6-8 μm for the DDB-MT data set, and 2-5 μm DDH-MT data set. Each tilt was acquired as movies in counting mode using a dose rate of 5.3 e^-^/pixel/s, with a per-frame exposure time of 80 ms and a dose of 0.09 e^-^/Å^2^. The total cumulative dose for each tilt series was 114 e^-^/Å^2^, and was distributed throughout the tilts based on the cosine of the tilt angle to account for changing sample thickness with increasing tilt. 154 and 126 tilt series were collected for DDB-MT and DDH-MT samples at a nominal magnification of 14,000X, giving a calibrated pixel size of 2.13 Å/pixel at the detector level.

Tilt series for the dynein-MT sample were collected on a Thermo Fisher Arctica electron microscope operating at 200 keV and equipped with a Gatan K2 Summit direct electron detector operating in movie mode, as described above. The total cumulative dose and dose distribution for each tilt series was same as described for DDB-MT and DDH-MT data sets. Data were collected using the Leginon package (Suloway 2005) with an alternating tilt scheme (Hagen 2017). A total of 58 tilt series were collected at a nominal magnification of 17,500X, giving a calibrated pixel size of 2.33 Å/pixel at the detector level.

### Tomogram reconstruction

Image processing and tomogram reconstructions were performed in similar fashion for all samples. Movie frames for each tilt were translationally aligned to account for beam-induced motion and drift using the GPU frame alignment program MotionCorr (Li 2013). A frame offset of 7 and a B-factor of 2000 pixels was used for frame alignment. The raw tilts were initially Fourier-binned by a factor of 2. All micrographs were aligned using the 5nm gold beads as fiducial markers, and further binned by a factor of 2 (final pixel size of 8.52 Å/pixel for DDB-MT and DDH-MT datasets, and 9.32 Å/pixel for dynein-MT dataset) for reconstruction in the IMOD package (Kremer 1996). Tomograms were reconstructed using simultaneous iterative reconstruction technique (SIRT) with seven iterations in IMOD provided sufficient contrast for the purposes of particle selection. Tomograms were also reconstructed by weighted back projection (WBP) for the purposes of subtomogram averaging.

### Subtomogram averaging and data processing

Sub-volumes containing DDB-MT, DDH-MT, or dynein-MT were manually picked from SIRT-reconstructed tomograms with the EMAN2 single-particle tomography boxer program (Galaz-Montoya 2015). Picked coordinates for each sub-volume were imported into the RELION 1.4 subtomogram averaging workflow (Bharat and Scheres 2016). 502 and 303 sub-volumes were extracted from the WBP reconstructions of the DDB-MT and DDH-MT datasets, respectively. Sub-volumes were extracted using a cube-size of 96 voxels for the DDB-MT and 84 voxels for the DDH-MT dataset. Reference-free 3D classification in RELION did not yield any structures resembling dynein or dynactin complexes, and instead predominantly produced averages of MTs. Attempts to remove signal from MTs by applying binary masks did not improve our ability to resolve the MT-bound complexes. To overcome this issue, we developed an assisted 3D subtomogram averaging procedure (Supplemental Figure 2), wherein we manually docked the available reconstruction of the dynein tail-dynactin-BICD2N (TBD) complex (EMDB 2860(Urnavicius 2015)) into the DDB-MT or DDH-MT sub-volumes using UCSF Chimera (Goddard 2007). The docked densities provided the rotational and translational parameters to generate initial subtomogram averages of the DDB and DDH complexes. These initial averages contained recognizable molecular features consistent with the previously published TDB structure (Urnavicius 2015) (Supplemental Figure 2b). To better resolve different components (dynein tail-adaptor-dynactin region and dynein motors) of the DDB and DDH complexes, focused 3D refinements were performed using 3D ellipsoidal binary masks corresponding to the individual sub-regions (Supplemental Figure 2c). For each component, 3D refinement was performed in RELION using the initial alignment parameters, with a HEALPix order of 3, an angular step size of 7.5º, and an offset range of 5 pixels. These refinements resulted in better-defined sub-regions of the MT-DDB and MT-DDH complexes (Supplemental Figure 2c). The final resolutions of these reconstructions are conservatively estimated to be ~38 Å (by Fourier Shell Correlation at a 0.5 cutoff) (Supplemental Figure 4, 6a). Composite reconstructions of the DDB-MT and DDH-MT complexes were generated by aligning and stitching together the focused reconstructions using the “vop maximum” function in UCSF Chimera (Goddard 2007), which retains the maximum voxel values of overlapping volumes.

Although the presence of an additional tail dimer and appearance of 4 dynein motors in the DDB subtomogram average, as well as the absence of GFP in the DDH reconstruction, all serve as internal controls that preclude the introduction of model bias into our refinement procedure, we performed additional control experiments to rule out this possibility. We first tested the ability of our sub-volumes to reproduce the well-resolved dynein tail-adaptor-dynactin region after focused refinement of the motor domains. Focused refinement of the motors results in misalignment of the dynein tail-adaptor-dynactin region, resulting in poorly-resolved dynactin density. Re-refining this region using an ellipsoidal binary mask reproduces the dynactin with well-resolved structural features. Next, we docked the TDB complex (EMDB 2860 (Urnavicius 2015)) into the sub-volumes using randomly assigned Euler angles and performed the same refinement strategy outlined above. This was repeated using three unique seeds for randomization, and in each case the resulting subtomogram did not yield a recognizable complex (Supplemental Figure 3b).

229 dynein-MT sub-volumes were extracted from the WBP tomograms with a cube-size of 96 voxels. As with the DDB and DDH datasets, ab-initio 3D classification mostly resulted in MT averages, and did not yield any recognizable dynein structures. We attempted to perform an assisted alignment approach, which involved placing spherical markers on the individual dynein motor domains using IMOD(Kremer 1996). However, due to the variability the inter-motor spacing and disordered arrangement of the dyneins relative to the MTs, we were unable to produce a 3D subtomogram average of MT-bound dimeric dynein. The spherical markers were used to measure the inter-motor distances shown in Supplemental Figure 7.

Available crystal structures and atomic models of individual components (see Supplemental Table 1) were fitted into the final reconstructions using UCSF Chimera (Goddard 2007).

**Table 1.** Atomic models used to generate the pseudo-atomic model of the DDB-MT complex.

| **Atomic model** | **PDB Identifier(s)** |
| --- | --- |
| Dynein motor domain—AMPPNP-bound AAA+ ATPase domain | 4W8F (Bhabha 2014) |
| Dynein motor domain—Stalk | 3VKG (Kon 2012) |
| Dynein motor domain—microtubule-binding domain (MTBD), high affinity state | 3J1T (Redwine 2012) |
| Human cytoplasmic dynein-1 heavy chain and associated subunits (IC, LIC, LC7) bound to Dynactin and N-terminal GFP-BICD2N | 5NW4, 5NVS (Zhang 2017) |
| GFP | 1GFL (Yang 1996) |
| Dynein-dynactin-GFP-BICD2N complex | 2AFU (Urnavicius 2015) |
| Dynein motor domain—C-terminal domain | 3VKH (Kon 2012) |

### Data deposition

Reconstructed maps of DDB-MT and DDH-MT were deposited in EM Data Bank with accession IDs EMD-7000 and EMD-7001, respectively.

---

**Supplemental Figure 1.**
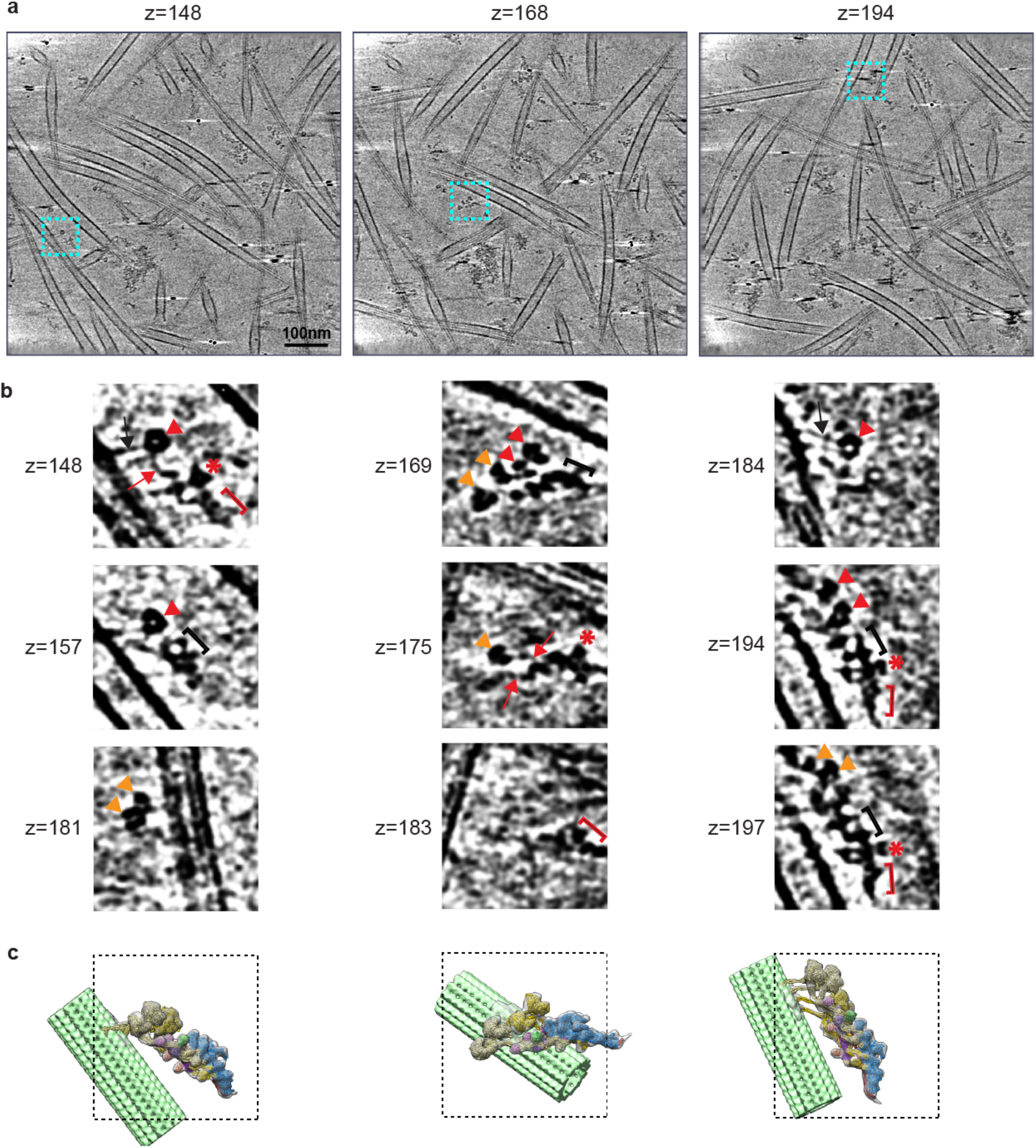
Representative cryo-electron tomographic reconstruction of MT-bound DDB complexes. **a**. Representative X-Y slices of a reconstructed tomogram progressing through the z-axis, displaying MTs with associated DDB complexes, and three representative complexes highlighted by cyan dashed squares. **b**. Representative X-Y slices progressing through the z-axis of extracted subvolumes demarcated by the colored dashed squares in **a**. These slices display several distinct features of DDB complexes, such as dynein dimer-1 motor domains (red arrow head), dynein dimer-2 motor domains (orange arrow head), dynein motor stalk (black arrow), dynein linker arm extension (red arrow), dynactin barbed end (black bracket), dynactin pointed end (red bracket) and dynactin shoulder domain (red asterisk). Scale bar in (a) represents 100 nm. **c**. Subtomogram average of the DDB-MT complex with fitted atomic models (as shown in Figure 1) oriented to match the DDB complexes shown in the corresponding extracted subvolumes, with a dotted box depicting the region shown above.

**Supplemental Figure 2.**
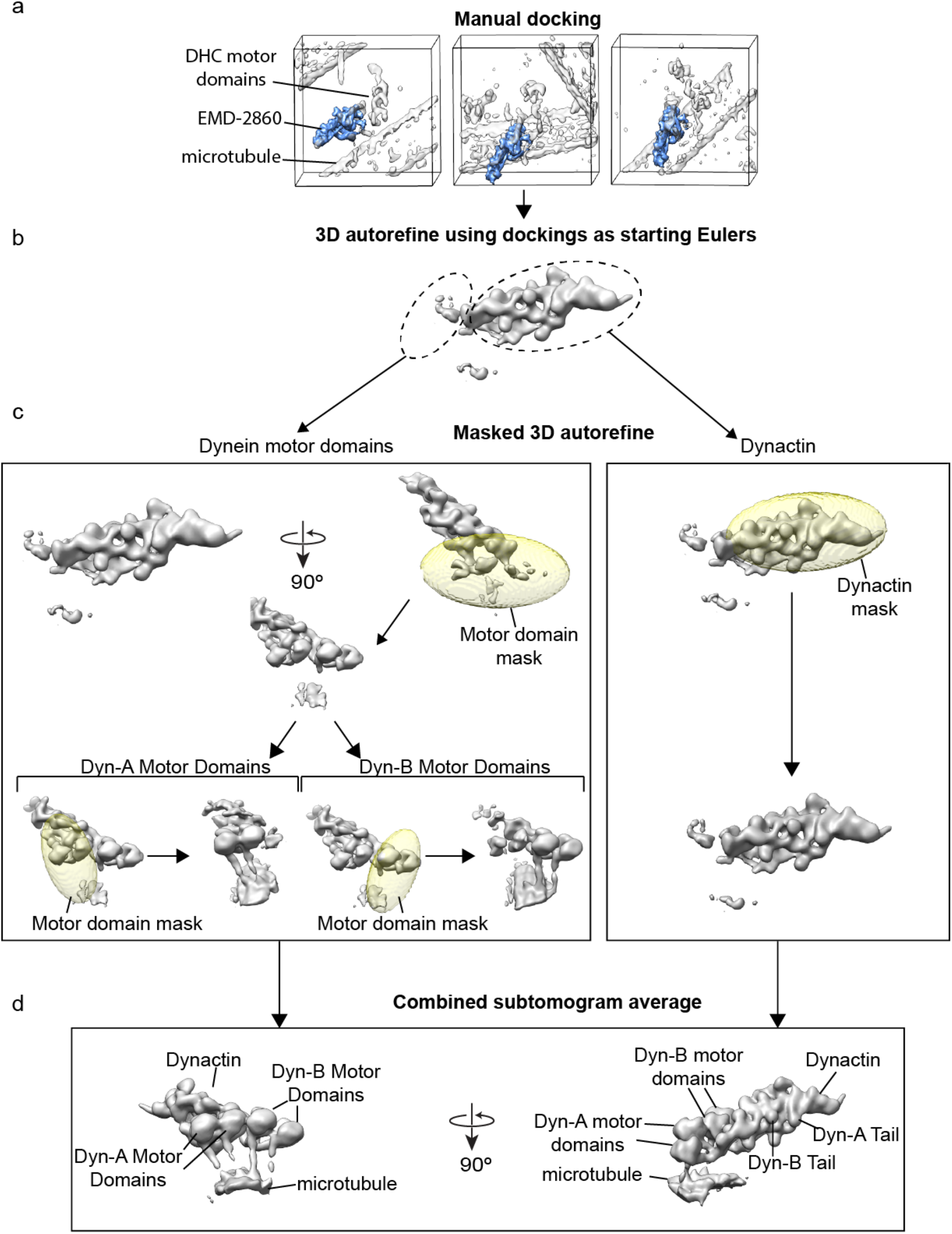
Subtomogram averaging and processing workflow. **a**. Cryo-EM reconstruction of the TBD complex (EMDB 2860 (Urnavicius 2015)) was docked into the DDB-MT and DDH-MT sub-volumes using UCSF Chimera (Goddard 2007). Selected subvolumes (gray density) are shown with docked TBD complex shown in blue. **b**. The docked densities provided the initial Euler parameters for RELION 3D autorefinement using a limited angular search, yielding subtomogram averages of the DDB and DDH complexes. **c**. Focused 3D refinements were performed using local 3D ellipsoidal binary masks (transparent pink) corresponding to dynein motor domains (left box) and dynein tail-adaptor-dynactin region (right box). In order to further resolve dimeric dynein heads belonging to each dynein (Dyn-A and Dyn-B), local masking and refinement was performed. **c**. Focused subvolume averages of different subregions of DDB-MT complex were aligned and stitched together using UCSF Chimera (Goddard 2007) to generate a composite map of the full complex. Orthogonal views of the composite map are shown.

**Supplemental Figure 3.**
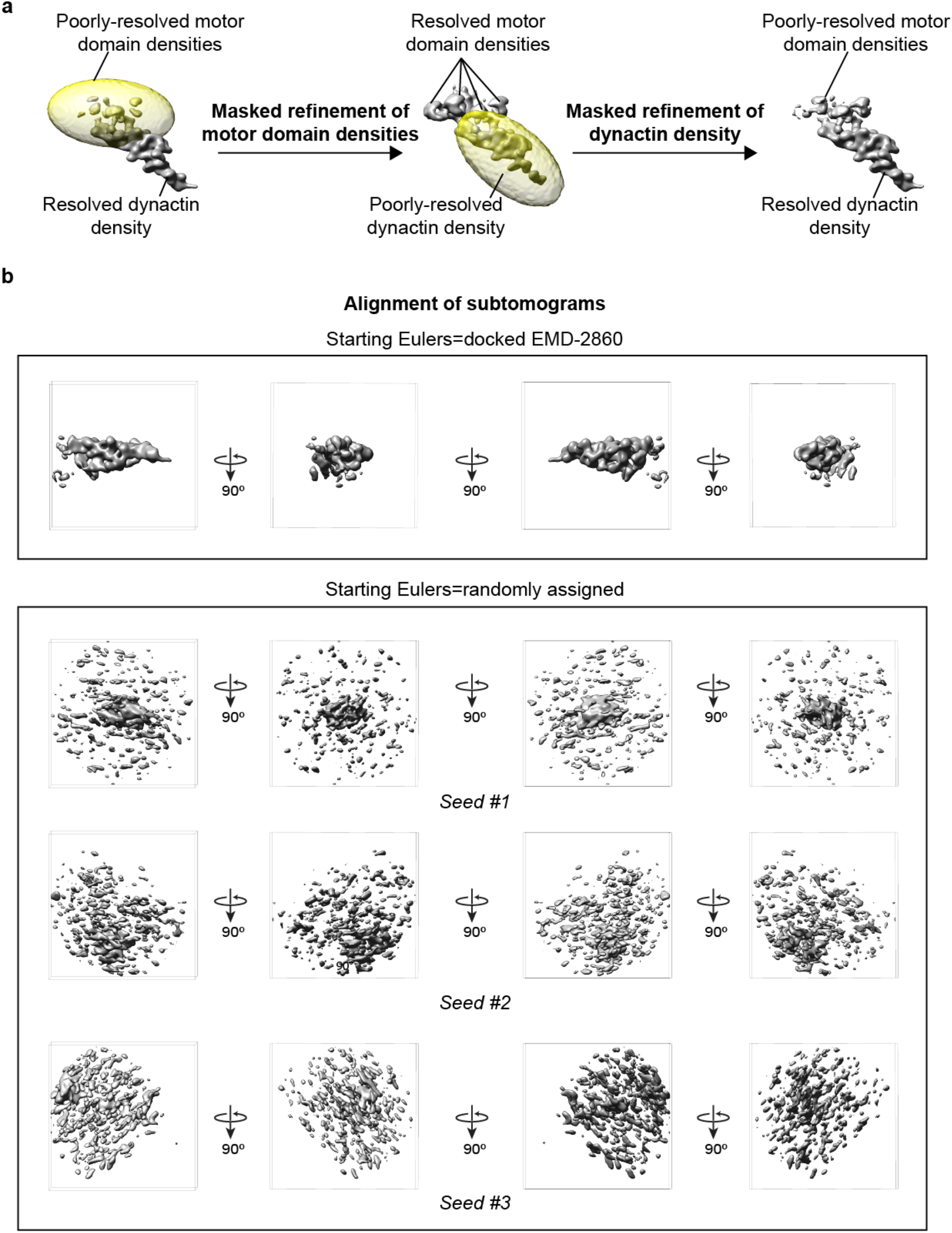
Validation of the subtomogram averaging procedure. **a**. 3D refinement focusing on the dynactin-dynein tail region of DDB-MT subvolumes results in this region becoming better resolved, while the density corresponding to the dynein motor domains worsens due to misalignment (left image). Re-refinement of the poorly-resolved dynein heads with ellipsoid binary mask (transparent yellow) results in better resolved density for the dynein motor domains, and poorly-resolved density dynactin-dynein tail (middle image). Continuing refinement with a binary mask around the dynactin-dynein tail density restores this density to a well-resolved state (right image), similar to that of the original 3D refinement of this region (left image), suggesting that the assisted subtomogram averaging procedure followed by focused 3D refinement does not impose any form of model bias to the final subtomogram averages. **b**. Assigning starting Eulers from the manually-docked TDB complex density (EMDB 2860 (Urnavicius 2015)) into the DDB-MT sub-volumes results in convergence to a density that corresponds to the dynactin-dynein tail-BICD2N region of the DDB-MT complex (top panel). Random assignment of the starting Eulers for the TDB complexes using three different seed models (Seed #1, #2, #3) results in 3D reconstructions that do not resemble any distinguishable feature of the DDB-MT complex, further suggesting that the assisted subtomogram averaging procedure does not introduce model bias to the final subtomogram averages of the dynein-dynactin-cargo adaptor complexes.

**Supplemental Figure 4.**
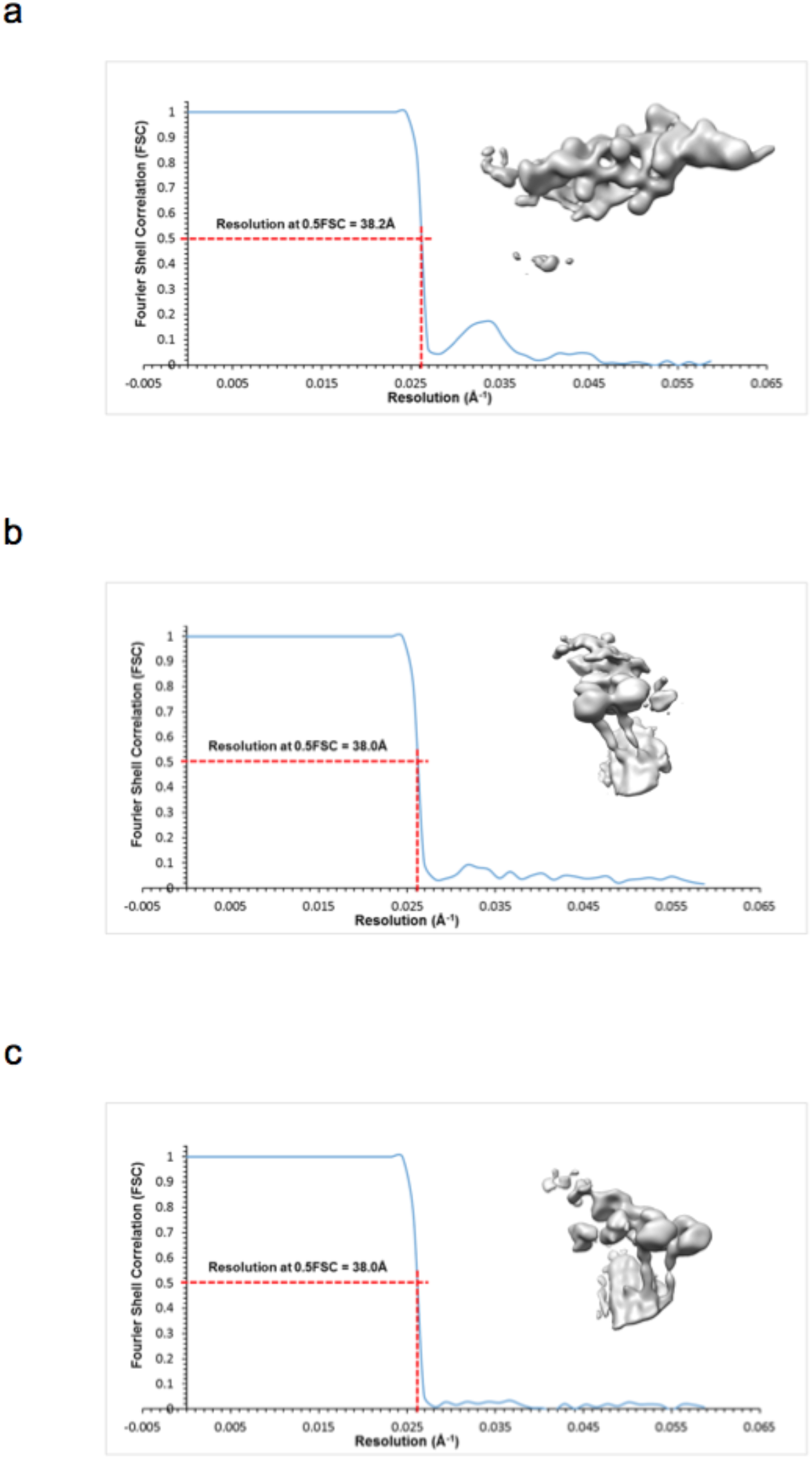
Focused subtomogram averages of DDB-MT complex were resolved to ~38Å resolution. **a-c**. Fourier shell correlation plots of individual focused reconstructions are shown with resolution reported at 0.5 FSC and the reconstructed maps shown to the right of the FSC curves. The dynactin-dynein tail region was resolved to 38Å (a), and Dyn-A (b) and Dyn-B (c) were both resolved to 38Å.

**Supplemental Figure 5.**
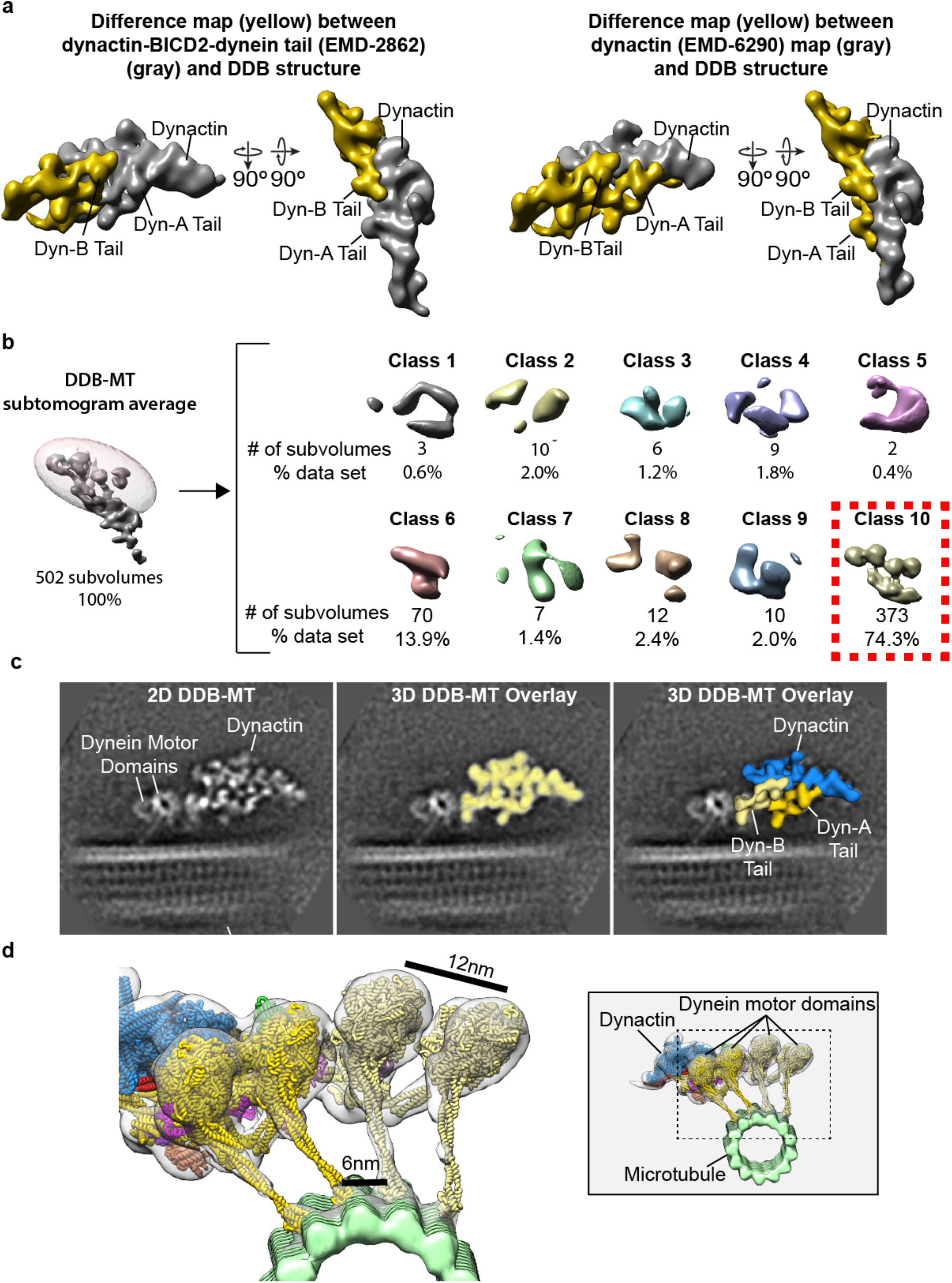
Two dimeric dyneins bind to a single dynactin in presence of cargo adaptor protein BICD2N. **a**. A difference map (gold), calculated by subtracting TDB density (EMD-2862 (Urnavicius 2015)) from focused subtomogram average of the dynactin-dynein tail-BICD2N region (gray), displays a density corresponding to a second dynein tail (left panel). A difference map (gold), calculated by subtracting dynactin density (EMD-6290 (Chowdhury 2015)) (gray) from focused subtomogram average of the dynactin-dynein tail-BICD2N region, displays a density corresponding to two dynein tails associated with dynactin (right panel). **b**. Focused 3D classification of the dynein motor domains of the DDB-MT complex using an ellipsoid binary mask (transparent pink) results in one well-resolved 3D class (Class 10), containing majority of the subvolumes (74.3%), and shows the presence of four dynein motor domains corresponding to two dimeric dyneins. **c**. Comparison of 2D averages of DDB-MT complex (Chowdhury 2015) (left image) with DDB-MT’s dynactin-dynein tail-BICD2N subvolume average (light yellow density), by overlaying the 3D density map onto the 2D average, shows a high correlation between the two (middle image). Segmentation of the 3D subvolume average shows the presence of two dynein tails (light yellow and gold) associated with a single dynactin (blue) in DDB-MT complex (right image). **d**. Subtomogram average (gray transparent density) of the DDB-MT complex with fitted atomic models (as shown in Figure 1) shows that spacing between ATPase rings and microtubule-binding domains (MTBDs) is ~12nm and ~6nm, respectively. The position of the motor domains relative to the entire DDB-MT complex is shown in the inset (right panel).

**Supplemental Figure 6.**
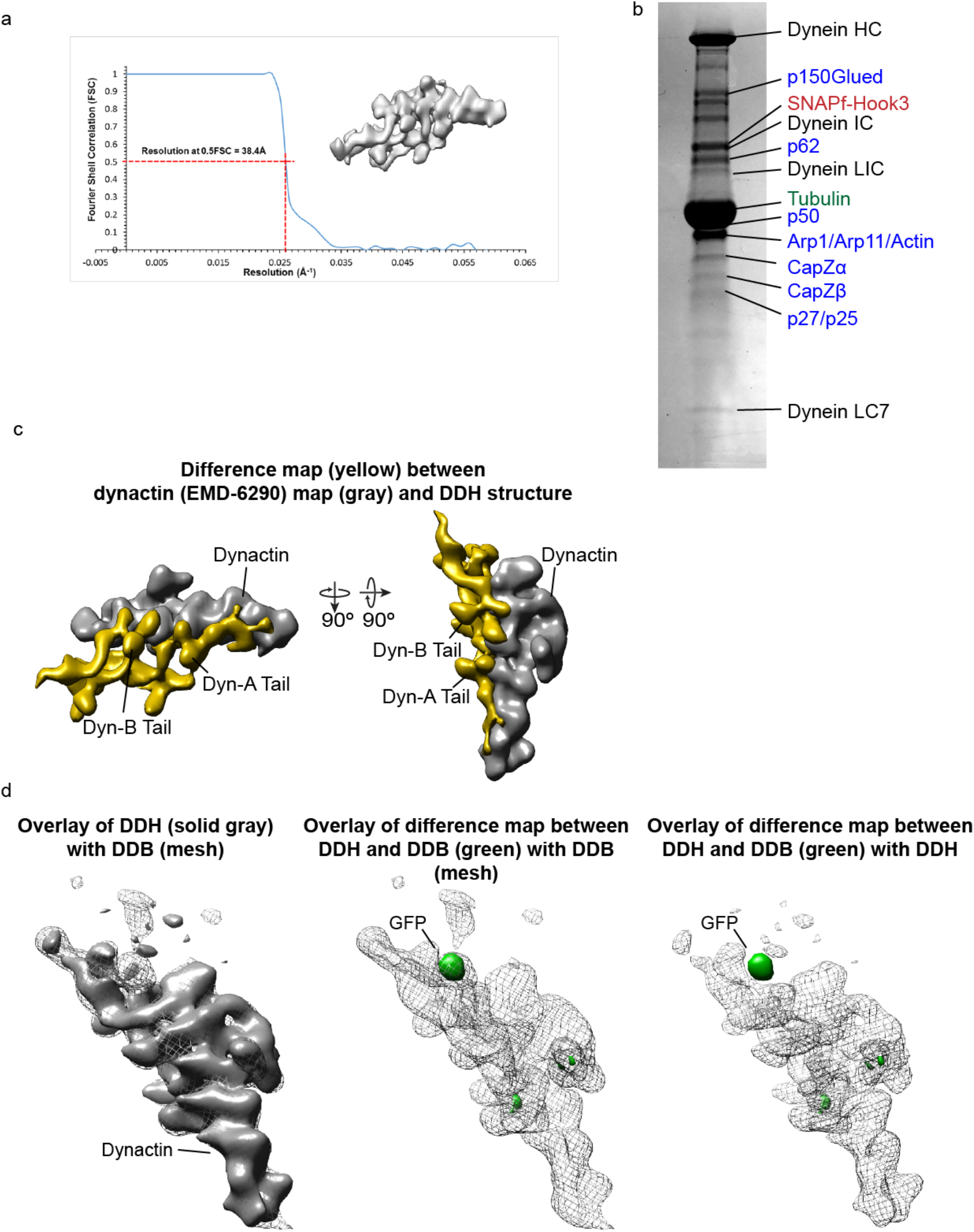
Subvolume average of DDH-MT complex. **a**. Focused subvolume average (map shown in gray) of the dynactin-dynein tails-Hook3 region of the DDH-MT complex was resolved to 38Å as shown in the FSC plot. **b**. SDS-Polyacrylamide gel electrophoresis (SDS-PAGE) of microtubule-bound dynein-dynactin-Hook3 complex purified from mouse brain shows that all components of the complex are present, including dynein and dynactin subunits, as well as Hook3 and tubulin. **c**. A Difference map (gold), calculated by subtracting dynactin density (EMD-6290 (Chowdhury 2015)) (gray map) from focused subtomogram average of the dynactin-dynein tail-Hook3 region, displays a density corresponding to two dynein tails associated with dynactin. **d**. The density for the dynactin-dynein tails-Hook3 complex (solid gray) correlates well with the corresponding density in the DDB-MT complex (mesh) (left image). A difference map (green), calculated by subtracting the DDH density from DDB, emphasizes a density corresponding to the N-terminal GFP tag of the BICD2N construct, which is present in the DDB-MT complex (middle image), but replaced with a smaller SNAPf tag in the Hook3 construct in the DDH-MT complex (right image).

**Supplemental Figure 7.**
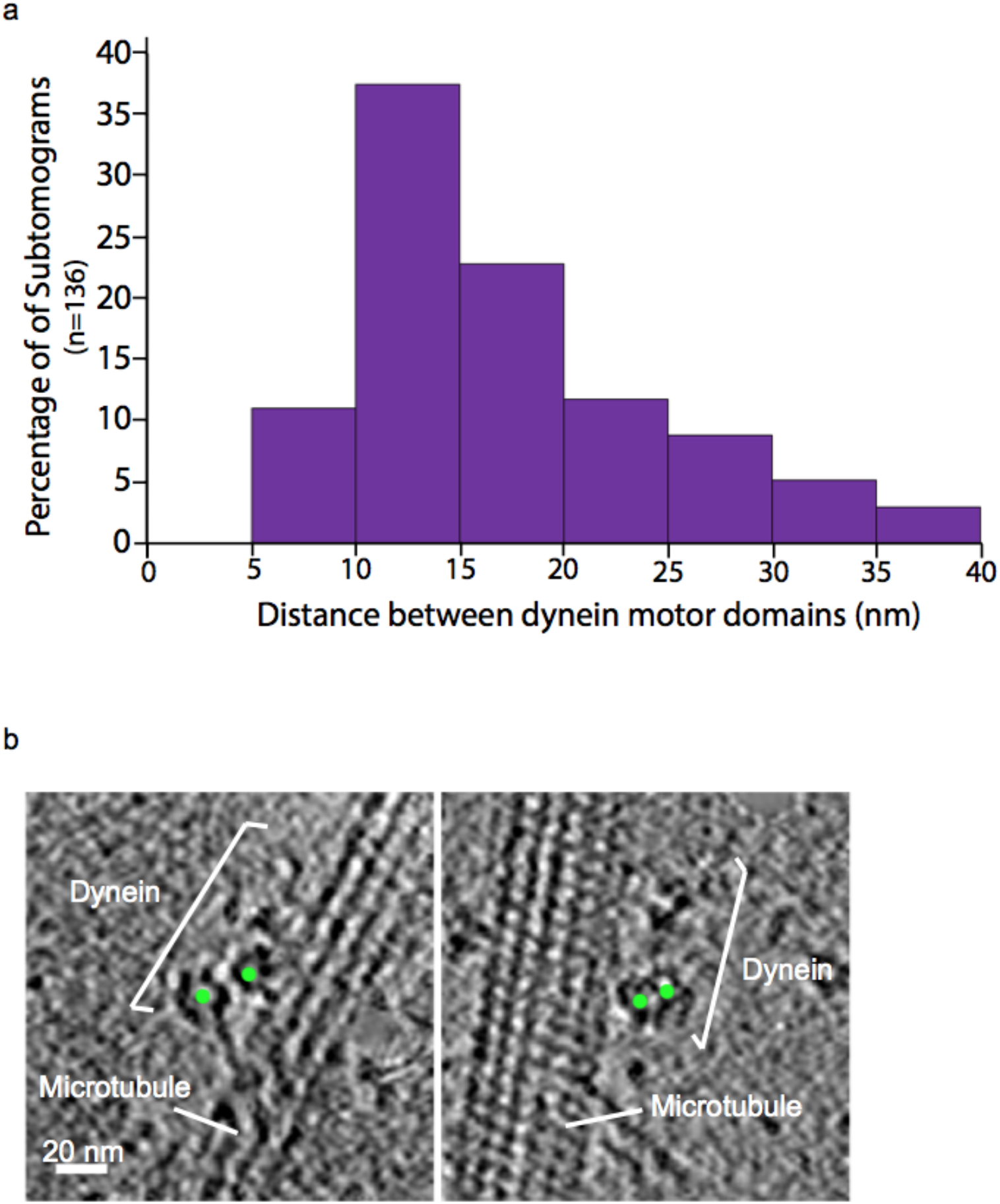
Dynein motor domains on MTs in the absence of cofactors. **a**. Histogram showing distribution of distances between motor domains of dimeric dyneins in DY-MT complexes indicates that the majority are spaced 10-15nm apart, however a wide range of distances is also observed (5-40nm), suggesting a high degree of flexibility of dimeric dynein in the absence of dynactin and cargo adaptors. Large distances (>25nm) may result from dyneins observed to be bridging neighboring MTs. **b**. Representative X-Y slices from extracted subvolumes of DY-MT complexes with the center of two motor domains marked with a green dot. Scale bar in (b) represents 20nm distance.

**Supplemental Figure 8.**
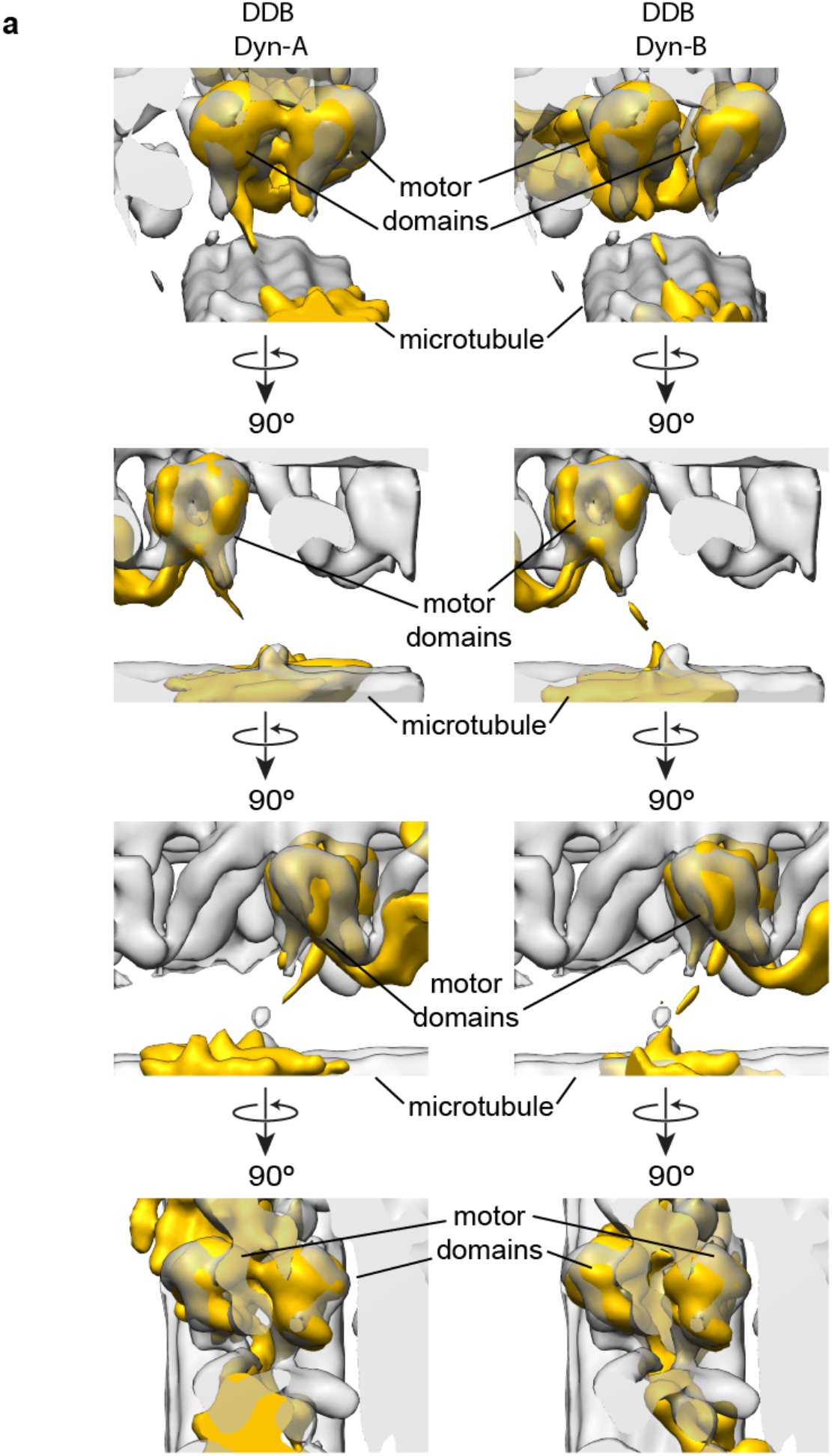
Similarities between the organization of cytoplasmic and axonemal dynein motor domains. **a**. Density corresponding to dynein motor domains with linker arm extensions from each of the dynactin-associated dynein dimers (Dyn-A, left column; Dyn-B, right column) (gold density) correlates well with the motor domains (gray density) present in the axonemal dynein map (EMD-5757 (Lin 2014)).

**Supplemental Figure 9.**
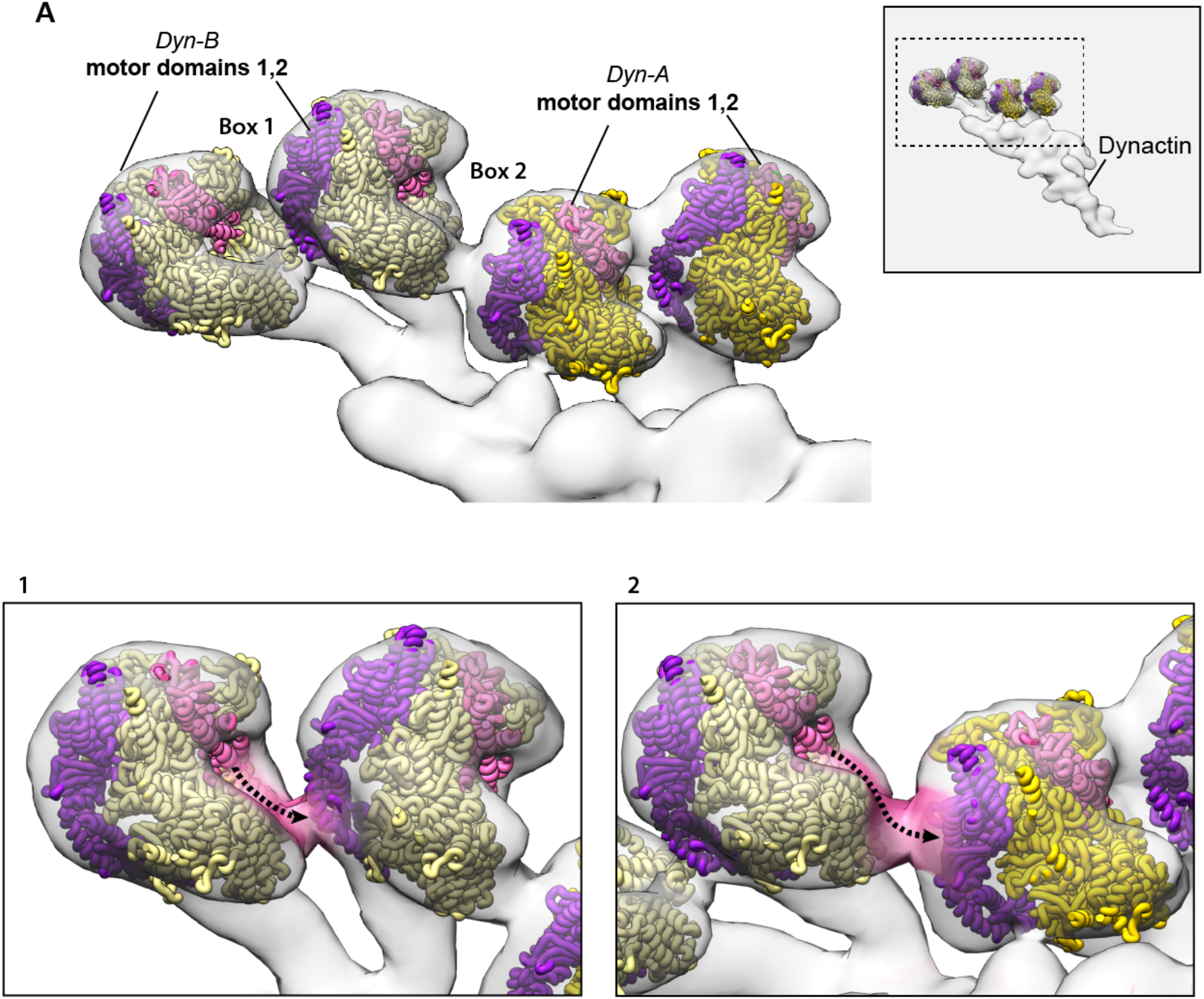
Putative intra - and inter-dynein motor domain contacts present in DDB-MT complex. **a**. Subtomogram average (gray transparent density) of the DDB-MT complex with fitted atomic models of dynein motor domains (yellow) shows putative intra-dimer (Box 1) and inter-dimer (Box 2) connections between adjacent motor domains. Box 1 and Box 2 in bottom panel have been magnified to show the connecting density (pink transparent density), which could be attributed to a proposed movement (dotted arrow) of the CT-cap (pink) for interaction with the linker arm (purple) of adjacent motor domains. This density may also correspond to auxiliary regulatory factors associated with the MD. The positioning of the motor domains relative to the DDB-MT complex is shown in the boxed region of the gray inset (upper right).

**Supplemental Movie 1: Subtomogram average of the dynein-dynactin-BICD2N complex** The subtomogram average is shown as a transparent gray density, and docked atomic models are colored as in Figure 1.

